# Growth characteristics of pneumococcus vary with the chemical composition of the capsule and with environmental conditions

**DOI:** 10.1101/416040

**Authors:** Adrienn Tothpal, Katherine Desobry, Shreyas Joshi, Anne L. Wyllie, Daniel M. Weinberger

**Author notes:** **Presented in part:** at ASM Microbe, New Orleans, 2017 and the 11th International Symposium on Pneumococci and Pneumococcal Diseases (ISPPD), Melbourne, 2018. **Corresponding authors**, Adrienn Tothpal: Mailing address: Institute of Medical Microbiology, Semmelweis University, Nagyvarad ter 4, Budapest, Hungary HU-1089. Daniel Martin Weinberger: Mailing address: Yale School of Public Health, 60 College Street, New Haven, CT 06510, USA.

## Abstract

**Background:** Pneumococcus, a bacterium that typically resides in the nasopharynx, is exposed to a variety of temperature and oxygen levels in the upper respiratory tract and as it invades the lung, tissues, and blood. The response to these variations likely varies by strain and could influence the fitness of a strain and its virulence. We sought to determine the effect of environmental variability on the growth characteristics of pneumococcus and to evaluate correlations between variability in growth characteristics between strains and biological and epidemiological characteristics.

**Methods:** We evaluated the effect of temperature and oxygen on the growth of 256 pneumococcal isolates representing 53 serotypes, recovered from healthy carriers and from disease patients. Strains were grown at a range of temperatures anaerobically or in ambient air with and without catalase and were monitored by reading the optical density. Regression models were used to evaluate bacterial and environmental factors associated with characteristics of the growth curves.

**Results:** Most isolates grew to the maximal density at the temperature of the nasopharynx (~33C) and under aerobic conditions (with catalase). Maximum density achieved was positively associated with the presence of N-acetylated sugars in the capsule and negatively associated with the presence of uronic acids. Reaching a greater density at an early time point was positively associated with the prevalence of serotypes among healthy carriers in the pre-vaccine period.

**Discussion:** Environmental variability affects the growth of pneumococcus, with notable differences between isolates and by serotype. Such variability could be influenced by characteristics of the capsule and might affect virulence and transmissibility.

---

*Streptococcus pneumoniae* (pneumococcus) is an opportunistic pathogen that resides in the human nasopharynx. The nasopharynx is considered to be the reservoir of transmission between individuals [1]. Pneumococci are diverse, with >90 serotypes (defined by the capsule polysaccharide) and has tremendous genetic variation, resulting from recombination. Serotypes vary in their prevalence among healthy carriers and in the likelihood that they will cause severe disease [2]. As conjugate vaccines against 7, 10, and 13 pneumococcal serotypes have been introduced, the vaccine-targeted serotypes have declined in frequency among healthy carriers and as causes of disease, while serotypes not targeted by the vaccine have increased in importance (serotype replacement). Next-generation conjugate vaccines are under development that target additional serotypes, and it is likely that these vaccines will lead to further serotype replacement. Understanding the factors that influence the fitness of these non-vaccine strains could help to anticipate future patterns of serotypes replacement and could aid in the design of more optimal vaccines.

The success of pneumococcus in the nasopharynx and the likelihood that it causes disease is likely driven, in part, by how it responds to variations in its local environment. In different anatomical sites within the human host, pneumococci are exposed to variable temperature and oxygen levels. In the nasopharynx, considered its main niche, the average temperature is around 33°C, with some differences between children and adults [3–6]. The core body temperature, which would be encountered during invasion into tissues, is 37°C. The temperature in the lungs is constantly changing based on the temperature of inhaled air but is generally lower than 37°C [7]. During infection by pneumococci or during viral co-infection (such as influenza or RSV), both external and internal temperature increases [8–12]. Oxygen levels also vary within the host. In the nasopharynx, bacteria on top of the mucus layer are exposed to almost ambient air (20% O_2_). Pneumococci in biofilms in the nasopharynx encounter lower levels of oxygen [13]. Entering the lower respiratory tract or the middle ear, pneumococci are exposed to micro-aerophilic conditions and to almost anaerobic conditions when present in blood and the cerebrospinal fluid (CSF) [14–17]. Likewise, mucus production during infection (i.e., due to influenza or RSV) can block the air passage and form micro-aerophilic (around 5% O_2_) or even anaerobic microenvironments [15, 16].

While pneumococci exist in a complex environment, simple *in vitro* experiments can be used to isolate the response to specific environmental conditions. Laboratory studies have evaluated variations in growth characteristics by serotype and have identified a relationship between *in vitro* growth characteristics and prevalence of serotypes in the nasopharynx of healthy children [18, 19]. However, the effect of variation in temperature and oxygen on the growth of different strains and serotypes has not been systematically explored.

The aim of this study was to investigate how environmental variability in temperature and oxygen influences the growth characteristics of pneumococci and how the responses to these variations correlates with the biological characteristics of the strains and the observed epidemiology of the serotypes. Using a diverse set of clinical and nasopharyngeal isolates, as well as capsule-switch and capsule-knockout variants generated in the lab, we quantified how the growth characteristics of pneumococci *in vitro* vary under a range of temperatures and in aerobic and anaerobic conditions. Using statistical models, we estimated the variation in these growth characteristics associated with serotype (after adjusting for isolate-specific effects that could reflect culture history or other characteristics) and evaluated the relationship between the serotype growth characteristics and relevant serotype-specific epidemiological and biological variables.

## Results

### Descriptive results

In total, we performed more than 4,900 growth curves on 256 different pneumococcal strains, representing 53 different serotypes **(Figure 1)**. Because of the relatively poor growth achieved in aerobic conditions without catalase, all further analyses presented here focus on the results from clinical samples grown either with catalase or anaerobically (N=3151). There was a weak, positive correlation (rho=0.27, 95%CI: 0.24, 0.30) between the maximum density achieved and maximum growth rate and no notable correlation between the length of the lag phase and either growth rate (−0.06; 95%CI: −0.09, −0.02) or maximum density (0.04; 95%CI: 0.00, 0.07). Growth curves for all isolates and conditions can be explored interactively at https://weinbergerlab.shinyapps.io/ShinyGrowth_v2/.

**Figure 1.**
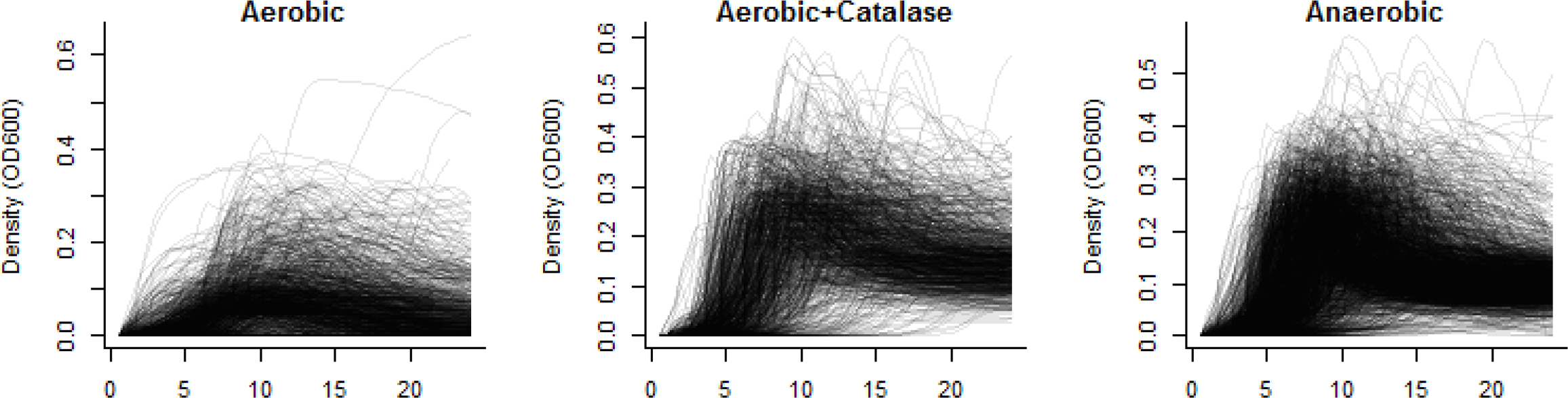
Variation in growth curves among all isolates used in this study. Growth was measured every 30 minutes over 24 hours at 30-39°C. (A) Ambient air, (B) ambient air with catalase, and (C) anaerobic conditions. Each line corresponds to an individual growth curve.

The maximum density of the strains was similar at 30-35°C, and lower densities were achieved at 37-39°C (**Figure 2A**); maximum density was greater in aerobic conditions (with catalase) than in anaerobic conditions. The growth rates were fastest at 35-37°C, with slower growth at lower or higher temperatures (**Figure 2B**); growth was faster in aerobic conditions (with catalase) than anaerobic conditions. The length of the lag phase increased with temperature (**Figure 2C**).

**Figure 2.**
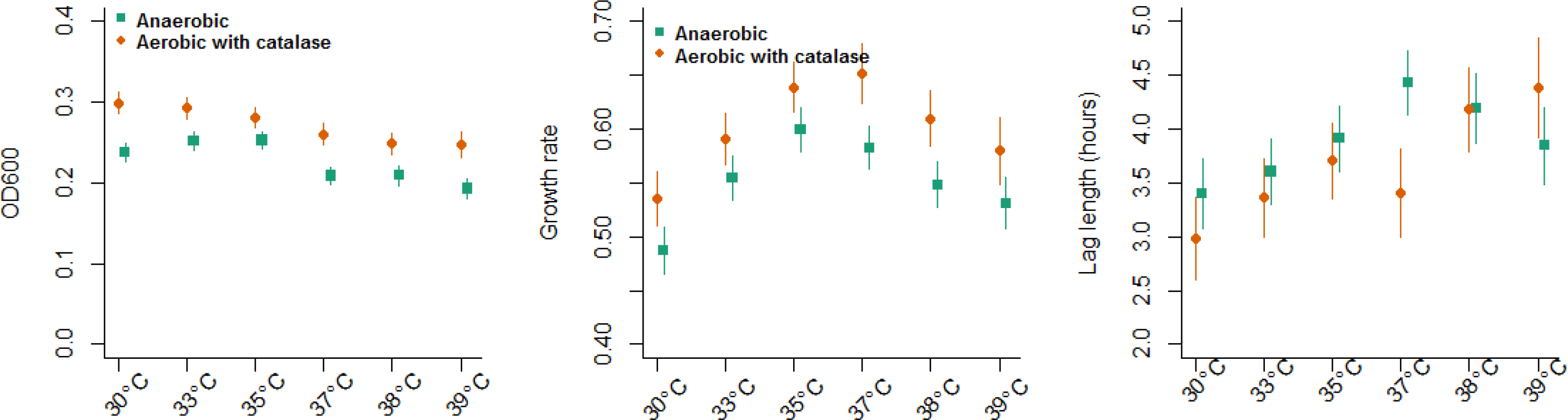
Effect of temperature and oxygen on growth phenotypes. **(A) Maximum density achieved, (B) Maximum growth rate and (C) length of lag phase.** Anaerobic (green squares) and aerobic+catalase (orange circles). These estimates are based on 3151 growth curves. Mean+/− 95% confidence intervals, calculated from a regression model adjusting for serotypes, presence of oxygen, temperature, site of isolation, and an interaction between temperature and presence of oxygen.

### Variation in growth characteristics associated with serotype

We quantified variation in maximum density achieved, growth rate, and length of lag phase associated with serotype. A number of serotypes differed from the reference (serotype 14) in maximum density achieved, length of the lag phase, and the density at an early time point (**Figure 3)**. We therefore evaluated characteristics between these serotype-specific averages and characteristics of the capsules. The presence of uronic acid (GlcA/GalA) in the capsular polysaccharide was associated with lower densities of growth and longer lag phases (P=0.001**, Figure 3)**. The presence of N-acetylated sugar in the capsule was associated with higher density (P=0.002), but not with growth rate or length of the lag phase (**Figure 4**). The strength of these effects did not differ appreciably between aerobic and anaerobic conditions.

**Figure 3.**
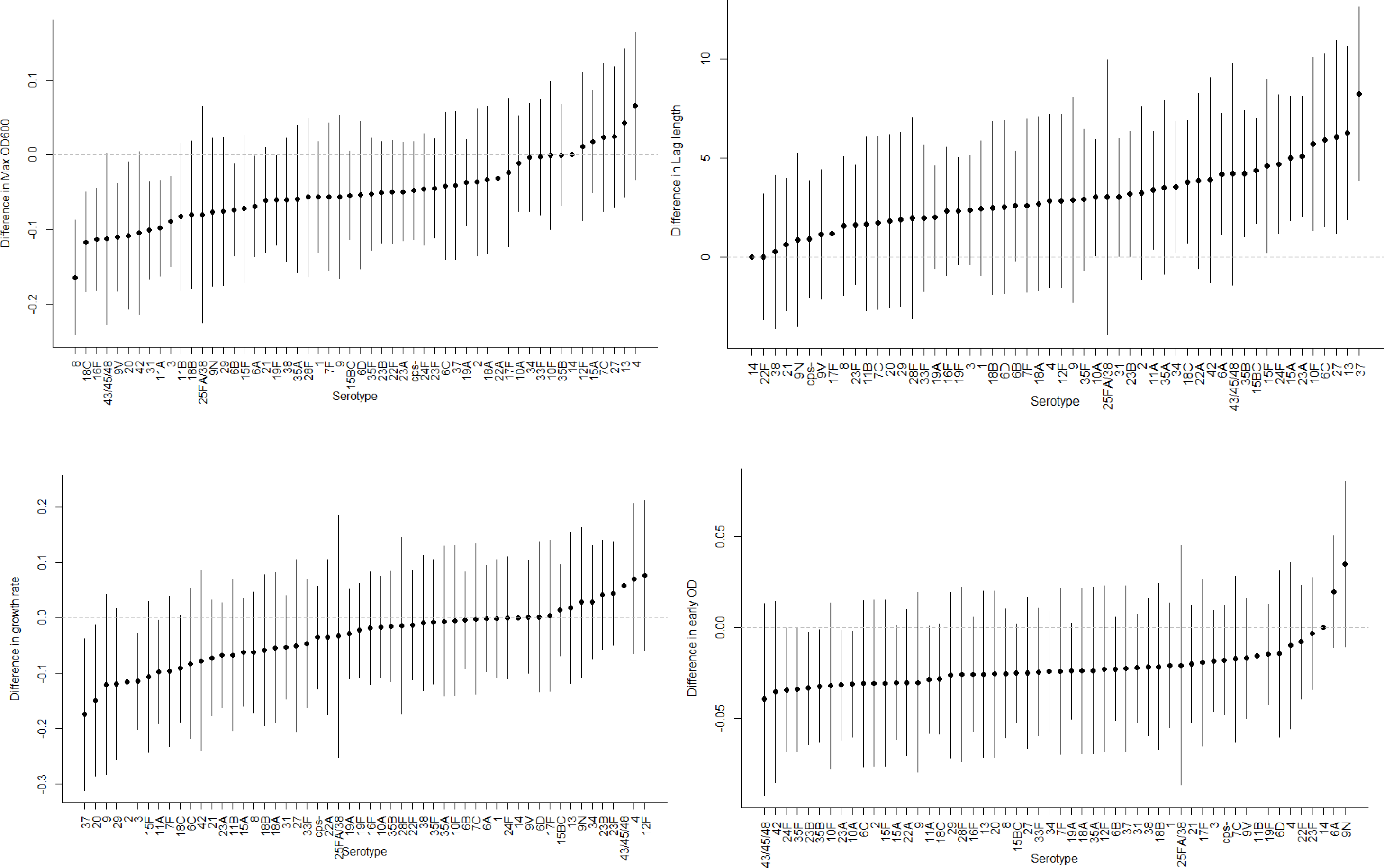
Variation in growth characteristics associated with serotype. Variation in (A) maximum density (B) length of the lag phase (C) maximum growth rates and (D) density at an early time point. The dots represent regression coefficients +/− 95% confidence intervals, as calculated from a regression model that controls for presence of oxygen, temperature, site of isolation, and interactions among these. The reference in the regressions is serotype 14.

**Figure 4.**
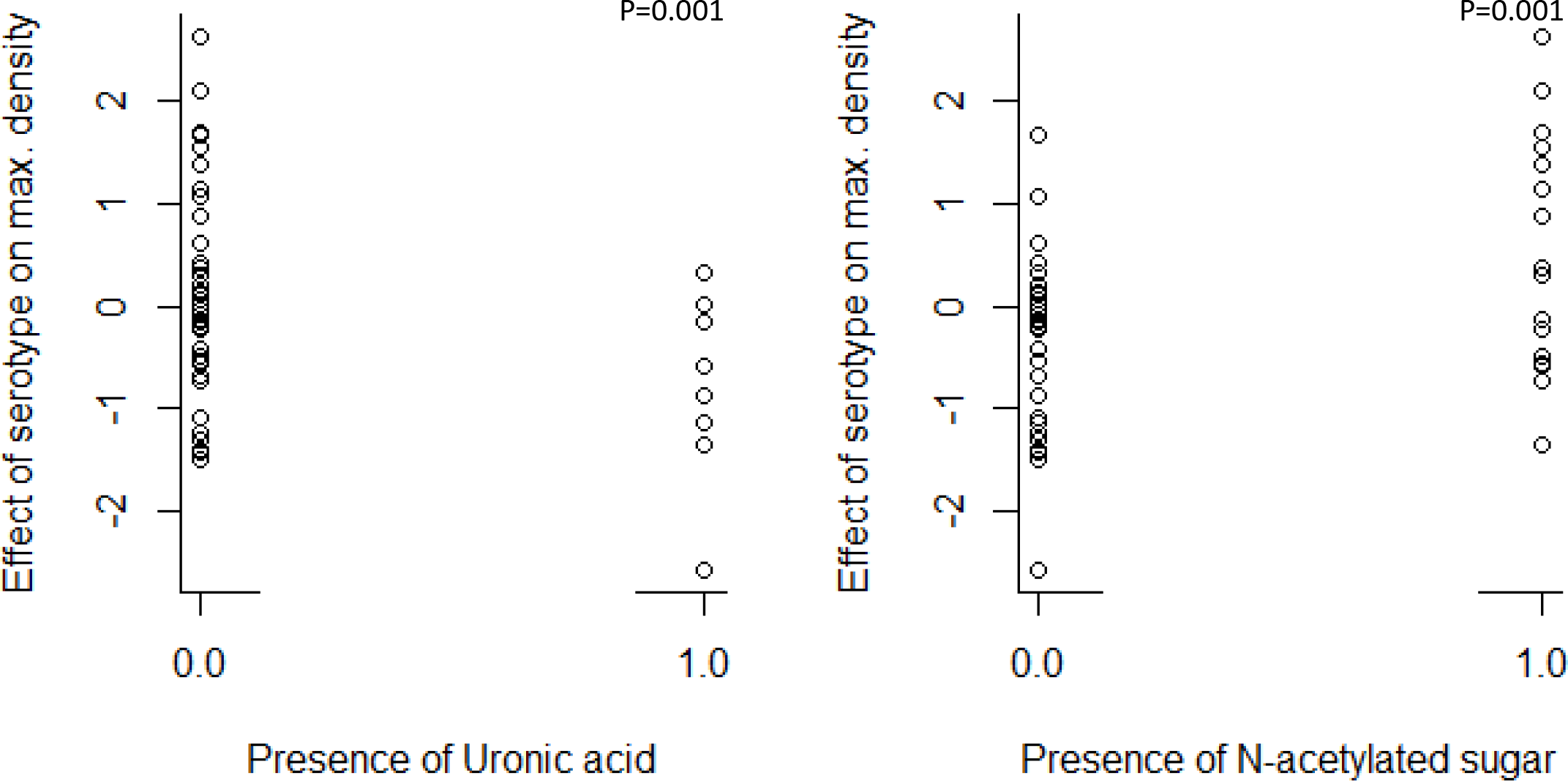
Variation in the maximum density achieved by serotypes based on characteristics of the capsule. The y-axis denotes the value of the serotype-specific regression coefficient, estimated for a model that adjusts for environmental conditions. The reference in the regression is serotype 14. Serotypes with GlcA/GalA in the capsule grew to a lower density on average, while serotypes with N-acetylated sugars in the capsule grew to a higher density. P-values are calculated with Monte Carlo resampling.

We next sought to determine whether this variability in growth characteristics was due to serotype or due to other genetic variability. Some of the serotypes in our collection were represented by multiple genetic lineages (MLST types). There did appear to be variation in growth characteristics associated with serotype that was similar across multiple isolates and MLST lineages (**Figures S1**, **S2**). However, there was not enough diversity in our sample to do a formal analysis. We also evaluated the growth characteristics of capsule-knockout strains as well as several capsule-switch variants. While the results were ambiguous, they suggested that there was an effect of capsule production on maximum density, and this effect was more pronounced during anaerobic growth compared with aerobic growth with catalase (**Figure S3**–**S6**).

### Effect of oxygen on growth varies by serotype

We considered whether the effect of oxygen on growth patterns varied by serotype (**Figure 5**). Overall, isolates producing certain capsules (e.g., 2, 4, 13, 35B) grew to a higher density under aerobic conditions (with catalase), while isolates producing other capsules (e.g., 6B, 8, 9N, 12F) did not show a difference between aerobic and anaerobic growth (**Figure 5**). This pattern was similar for maximum growth rate. For lag phase, there was little difference among the serotypes in how they responded to the presence of oxygen, with only the serotype 6C and 13 isolates exhibiting a longer lag phase in aerobic conditions.

**Figure 5.**
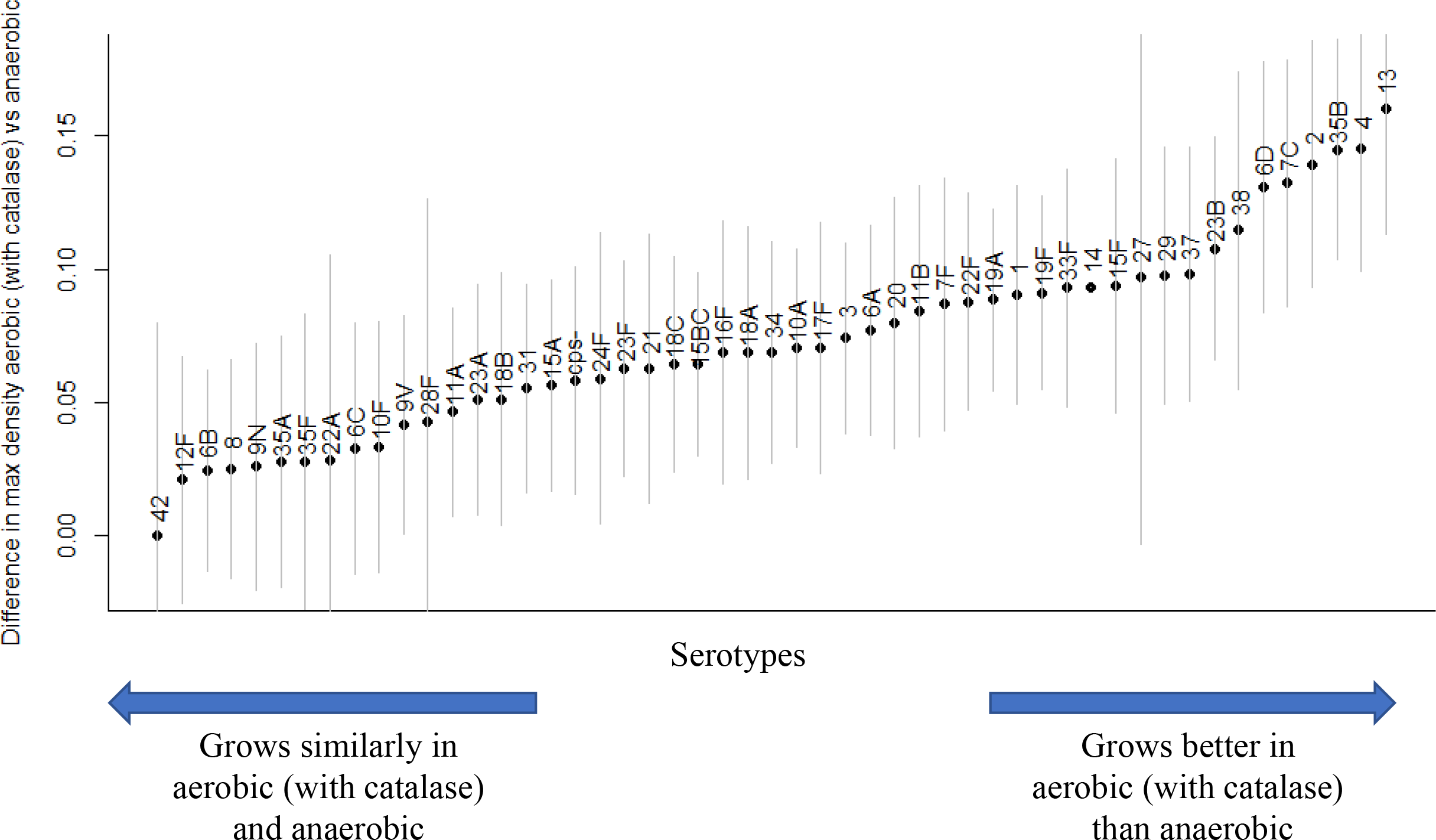
Difference in maximum density by serotypes when grown in aerobic conditions with catalase versus anaerobic conditions. Positive values indicate that a higher density (as measured by optical density) was achieved under aerobic conditions. A value of 0 indicates no difference. Mean+/− 95% confidence interval, as calculated using a regression model with covariates for serotype, presence of oxygen, temperature, an interaction between serotype and presence of oxygen, and a random intercept for the isolate. The reference in the regression is serotype 14.

### Relationship of growth characteristics to serotype-specific characteristics

Finally, we evaluated associations between the characteristics of the growth curves and serotype-specific epidemiological characteristics. There was a correlation between the density of a serotype at an early time point and the prevalence of the serotype among healthy carriers during the pre-vaccination period in several settings (**Figure 6**). The association was marginally stronger when looking at aerobic growth compared to anaerobic growth. There was no association between prevalence in carriage and maximum density achieved, length of the lag phase, or maximum growth rate. There was also no notable association between these growth characteristics and the invasiveness of the serotypes or the case fatality ratio.

**Figure 6.**
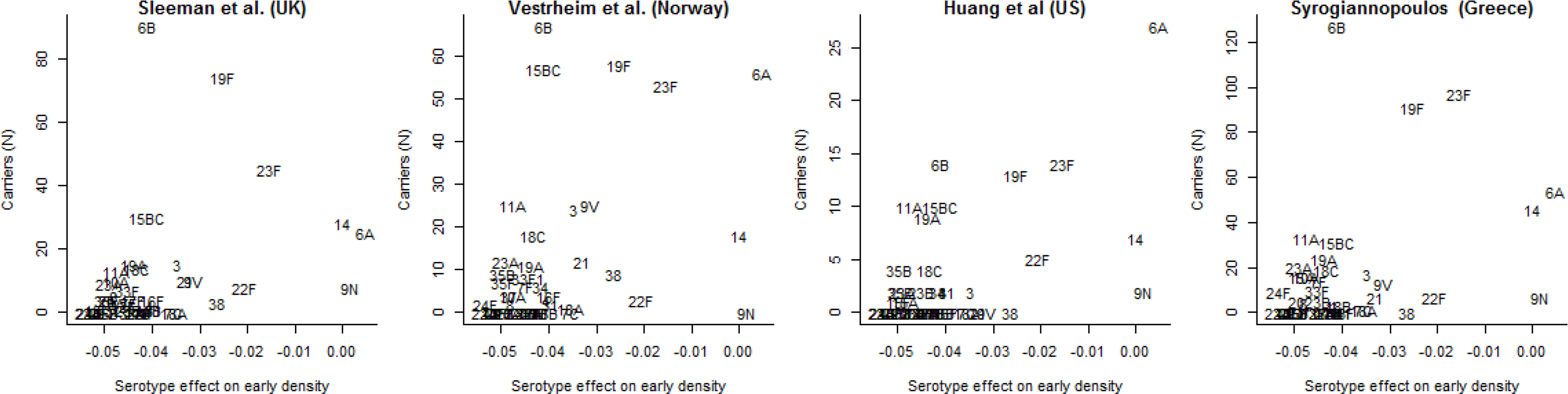
Correlation between carriage prevalence in the pre-vaccine period in different locations and the density of growth at an early time point. The serotype-specific estimates of the density of growth at an early time point are calculated from a regression model that controls for environmental variation and site of isolation. The Y-axis shows the number of carriers of a serotype, and the X axis shows the serotype-specific regression coefficient, which represents the variation in density for that serotype compared to serotype 14, after adjusting for environmental variables. The reference in the regressions is serotype 14.

## Discussion

We provide novel information about how the response to environmental conditions can differ between pneumococcal isolates. Testing a diverse set of strains, pneumococci grew to the highest density under conditions that mimic the normal environment of the nasopharynx in terms of temperature and oxygen level. Important differences in these patterns were observed between isolates. While, some of this variability could be due to the culturing history of the individual isolates, patterns of variation were apparent by serotype. The presence of specific polysaccharide components in the capsule was associated with higher or lower density. And the density of growth at an early time point was correlated with higher prevalence of those serotypes among healthy carriers.

Serotype is a major determinant of pneumococcal epidemiology and biology. We identified a novel association between the capsule composition and the growth phenotype of the isolates, with serotypes containing uronic acids in the capsule growing to a lower density and serotypes with N-acetylated sugars growing to a higher density. Further work is needed to understand the mechanism underlying these patterns, but the reliance on metabolic pathways shared between central metabolism and capsule production could influence the growth phenotypes. This could result from metabolic fluxes that restrict growth or through feedback loops that lead to increased metabolic activity. When comparing growth in aerobic conditions (with catalase) with growth in anaerobic conditions, the benefit of oxygen varied by serotype (**Figure 4**, **5**). Serotypes 2, 4, 13, 23B, 35B and 38 grew better with additional catalase than in anaerobic conditions, whereas serotypes 6B, 8, 9N, 12F, 22F and 42 grew similarly in both environments. These patterns did not correlate with capsule characteristics. We tested a relatively small number of isolates per serotype, so it is possible that non-capsular genetic variations or differences in the culture history of the isolates could influence the observed responses to oxygen.

As the nasopharynx is the normal habitat of pneumococcus, we had hypothesized that pneumococci would grow optimally at temperatures in the low-30C range, similar to the temperature of the nasopharynx. Indeed, temperature played an important role in terms of the maximal density achieved and how quickly the isolates started growing (lag phase). Isolates grew to the highest density and had the shortest lag at temperatures resembling those of the nasopharynx; these patterns were particularly pronounced for those isolates obtained from the nasopharynx. These patterns among the nasopharyngeal isolates are unlikely to be a result of the *in vitro* culture history of the strains alone—pneumococcal isolates are typically cultivated at 37°C, and thus if the patterns were a result of adaptation, we would expect more efficient growth at that temperature rather than at 33°C. These findings, along with recent work on the effect of lower temperatures on the immune response to pathogens in the upper respiratory tract [20], suggest that the environment of the nasopharynx is optimal for pneumococcal growth. IPD isolates responded differently to environmental variations.

During the invasion process, when the pneumococcus transitions from the nasopharyngeal environment to the internal body environment, it has to adapt to many changes, including nutrient, temperature and oxygen levels. Temperatures vary from the low-30°C range in the upper respiratory tract to 37°C in core body sites and even higher during fever. Likewise, oxygen levels can vary considerably during infection. Increased mucus production (due to co-infection with viruses or other pathogens in the upper respiratory tract) leads to lower oxygen levels and may generate a local hypoxic environment [21–23]. The availability of oxygen is also decreased in pneumonia, empyema, and otitis media. Oxygen levels in the uninflamed middle ear space, for example, resemble that of venous blood, are less than a third that of the airway, and may be further reduced by the presence of effusion [14, 15, 24]. Some isolates appeared to be strongly influenced by these variations in oxygen and temperature, indicating that the strains have the capability of adapting (either in the host or during *in vitro* growth) to thrive under different conditions.

Some of the differences observed in growth phenotypes between carriage and disease isolates could reflect opaque/transparent phase variation [25]. Opaque variants are generally isolated from IPD and have increased capsule production and decreased production of certain surface proteins. Phenotypically, the presence of oxygen accentuates differences in capsule production between opaque and transparent variants [24]. This effect could be mediated via the pathways involved in converting pyruvate to acetyl-CoA, an important biochemical precursor for capsule production for many serotypes [26]. Variations in the use of this pathway between serotypes or the efficiency of this pathway between lineages could influence some of the patterns that were observed in the growth curves.

This study had certain limitations. For the growth curves, we used BHI broth which is an artificial growth medium that differs in nutrient composition from the host. We evaluated several minimal media but found that growth was generally poor, making comparisons between strains difficult. While we tested a large number of strains representing many serotypes, some serotypes were only represented by a single isolate (i.e., 11B, 12F, 13). This could limit the generalizability of serotype-specific findings in these instances, making it difficult to make inferences about whether variability was due to serotype, site of isolation, or lineage effects. The strains used in this study were largely a convenience sample from clinical studies. The genetic diversity of pneumococcus makes it difficult to draw conclusions about the cause of differences between strains. The growth curves with the capsule-knockout strains and capsule-switch variants suggests that the capsule itself could influence these phenotypes. We did not perform any gene expression studies, which could be highly influenced by environmental conditions [27]. Further work could explore the genetic basis (both capsular and non-capsular factors) for the differences in growth phenotypes between strains.

In conclusion, we demonstrate that the growth characteristics of pneumococcus are influenced by environmental variations, that the effect of these variations depend on strains, and that the optimal growth conditions for carriage isolates resemble the conditions of the nasopharynx. Moreover, we demonstrate that the growth patterns among serotypes are associated with carriage prevalence and other epidemiological and biological characteristics. These results could help in understanding which of the serotypes has the greatest capacity to emerge - in both carriage and disease - in the future.

## Methods

### Bacterial strains, culture media, and chemicals

#### Strains

Invasive pneumococcal disease (IPD) isolates were obtained from the isolate bank at the Centers for Disease Control/Active Bacterial Core surveillance; carriage isolates were provided by Ron Dagan (Ben-Gurion University, Israel), Adrienn Tothpal and Eszter Kovacs (Semmelweis University, Hungary [28, 29]) and Debby Bogaert and Anne Wyllie (UMC, Utrecht [30]) (**Table 1**). Capsule-switch variants generated on the TIGR4 genetic background and the serotype 6B knockout strain were provided by Marc Lipsitch and generated as previously described [31]. Additional capsule-knockout strains were generated by replacement of the capsule biosynthesis locus with the Sweet Janus cassette [32].

**Table 1.** Pneumococcal strains used in this study.

| Source of isolate | Number of isolates | Country | Serotypes |
| --- | --- | --- | --- |
| Invasive pneumococcal disease (sepsis, meningitis) | 40 | US (CDC) | 1, 2, 3, 4, 6A, 6B, 6C, 6D, 7C, 7F, 8, 9N, 9V, 10A, 10F, 11A, 11B, 12F, 13, 14, 15A, 15B, 15C, 15F, 17F, 18A, 18B, 18C, 19A, 19B, 20, 22F, 23F, 29, 31, 33F, 34, 35A, 35B, 37 |
|  | 2 | Hungary | 3, 6B/D |
| Pneumonia | 8 | Hungary | 3, 8, 10A, 15A, 19F, 35B, 43/45/38, NT |
| Conjunctivitis | 16 |  | 3, 21, 31, 34, 42, 11A, 15A, 15B, 16F, 19A, 19F, 23A, NT |
| Carriage | 52 | Israel | 6B, 14, 15B/C, 19A, 19F, 23F, NT |
|  | 87 | Hungary | 1, 3, 8, 21, 31, 34, 38, 6A, 9V, 10A, 11A, 15A, 15B/C, 16F, 18C, 19A, 19F, 22A, 22F, 23A, 23B, 23F, 24F, 28F, 35F |
|  | 51 | Netherlands | 3, 27, 10A, 11A, 15B/C, 16F, 19A, 19F, 22A, 22F, 23B, 33F, 35B, 35F, NT |
| Laboratory-generated genetic variants | 4 | Various | TIGR4 (cps-, 5, 14, 19F), 603 (6B, cps-), CDC-10A (10A, cps-), CDC-15B (15B, cps-) |

The multi-locus sequence type was inferred for a subset of the isolates. These strains were subjected to Illumina NovaSeq sequencing [33], and MLST was determined using SRST2 [34].

#### Culture media

Pneumococcal isolates were stored at −80 °C on Cryobeads (Cryobank, Copan Diagnostics, Murrieta, CA). Strains were routinely grown at 37°C and 5% CO_2_ overnight on tryptic soy agar plates supplemented with 5% sheep blood (TSAII) (Thermo Fisher Scientific). Growth in broth culture was performed in BHI (Becton, Dickinson, and Co., Sparks, MD) with and without catalase (5000 units, Worthington Biochemical Corporation, Lakewood, NJ) for aerobic cultivation and with Oxyrase^®^ (Oxyrase, Inc., West Mansfield, OH) diluted 1:10 to create an anaerobic environment.

### Growth experiment

Strains were streaked onto TSAII plates and incubated at 37°C in a 5% CO2-enriched atmosphere overnight, then harvested into PBS to OD600 0.2 and diluted (6 fold) in BHI with or without catalase or Oxyrase. Growth was monitored in sterile flat-bottomed 96-well microtiter plates (BRAND GMBH, Wertheim, Germany) for 24 hours in a microplate reader (BioTek ELx808) with a built-in incubator, reading the optical density at 600 nm every 30 minutes, after 5s shaking (Gen5 program). Each strain was tested in all three oxygen conditions and across the full range of temperatures (30-39°C). An anaerobic control strain (*Bacteroides thetaiotaomicron*) was used to confirm the elimination of oxygen by Oxyrase.

### Data analysis

Each growth curve was blanked by subtracting the OD600 reading at t=0 for that well. In instances where the t=30 minutes measurement was lower than the t=0 measurement due to measurement error at the first time point, the OD600 at t=30m was subtracted instead.

We extracted three characteristics from each of the growth curves: maximum density achieved, length of the lag phase, and density achieved at an early time point. The length of the lag phase was determined by fitting a smoothing spline to the log-transformed growth curves (smooth.spline function in R, with a smoothing parameter of 0.5) and calculating the second derivative of this curve. The maximum of the second derivative gives the point at which the growth rate increases the most, corresponding to the transition from stationary phase to log phase. We were also interested in evaluating the density at an early time point. However, what constitutes an ‘early’ time point likely varies by environmental conditions and source of the isolates. Therefore, we determined the mean time when log-phase growth began for each temperature/oxygen/site of isolate combination, and then determined the OD600 for the corresponding isolates 1 hour after this point.

To quantify variations in the growth characteristics by serotype, site of isolation, and environmental condition while adjusting for repeated measurements and strain-to-strain variations, we used linear mixed effects models (lme4 package in R) [35]. The outcome variable was maximum OD600, fixed effects variables included serotype, temperature (categorical), oxygen (aerobic+catalase, aerobic without catalase, anaerobic), site of isolation (categorical), and a random intercept for each isolate. Certain interactions among the fixed effects were also evaluated to test specific hypotheses (site of isolate*oxygen; oxygen*temperature; serotype*oxygen). The significance of these interactions at different levels was evaluated using the interactionMeans function in the phia package in R [36].

#### Linkage to serotype-specific characteristics

We sought to evaluate the link between serotype-specific growth curve characteristics and previously published serotype-specific characteristics (capsule structure, disease severity, invasiveness, pre-vaccine carriage prevalence) [37–40]. We first fit a linear mixed effects model as described above that had dummy variables for serotype (reference: serotype 14) and controlled for temperature (categorical), presence of oxygen, site of isolation, and interactions between site of isolation and presence of oxygen and temperature and presence of oxygen. We then extracted the serotype-specific regression coefficients, which represent variation in the growth characteristics compared to serotype 14. These coefficients were used in second stage analyses to assess correlations with serotype-specific biological and epidemiological variables.

Associations with the presence of uronic acids or N-acetylated sugars were assessed using a Monte Carlo approach, where the mean maximum density was compared in the serotypes with or without the sugar components. The sugar component labels were randomly scrambled 999 times, and the difference was calculated for each of these samples. P-values were calculated by comparing the observed difference in means with the distribution obtained from the resampling. The association with carriage prevalence was assessed with Poisson regression. In a series of linear regressions, we evaluated the association case-fatality rate, complexity of the capsular polysaccharide (carbons/polysaccharide repeat) and invasiveness (log transformed).

#### Data availability

The raw data and an R Markdown file are available in a github repository (https://github.com/weinbergerlab/GrowthVariation) and can be used to fully re-create the analyses presented here.

## Acknowledgements

The authors gratefully acknowledge Marc Lipsitch, Krzysztof Trzcinski and Claudette Thompson for providing the TIGR4 and serotype 6B capsule-switch variants, the isolate bank from the Active Bacterial Core surveillance (ABCs)/Emerging Infections Programs (EIP) Network for providing the IPD isolates from the United States, Ron Dagan for providing the isolates from Israel, and Eszter Kovacs and Orsolya Dobay for providing the isolates from Hungary.

---

**Supplementary Figure 1.**
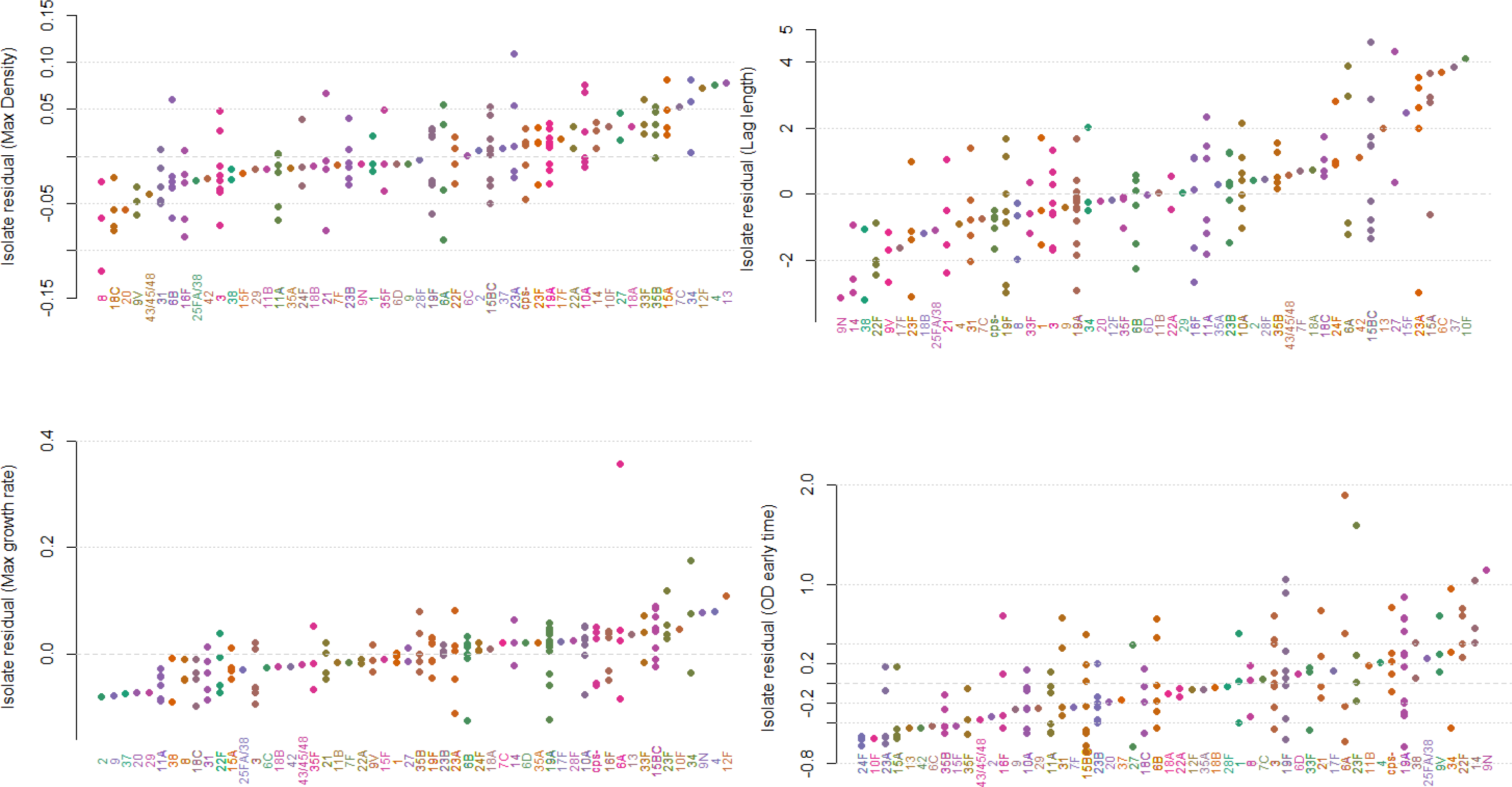
Isolate-specific random effects, organized by serotype. The random effect indicates how far above or below a strain is in maximum density achieved, lag length, growth rate, or density at an early time point. Serotype is not adjusted for in this model. The random effects are effectively an average residual for the isolate, after adjusting for temperature, site of isolation, and presence of oxygen. Each dot represents an individual isolate.

**Supplementary Figure 2.**
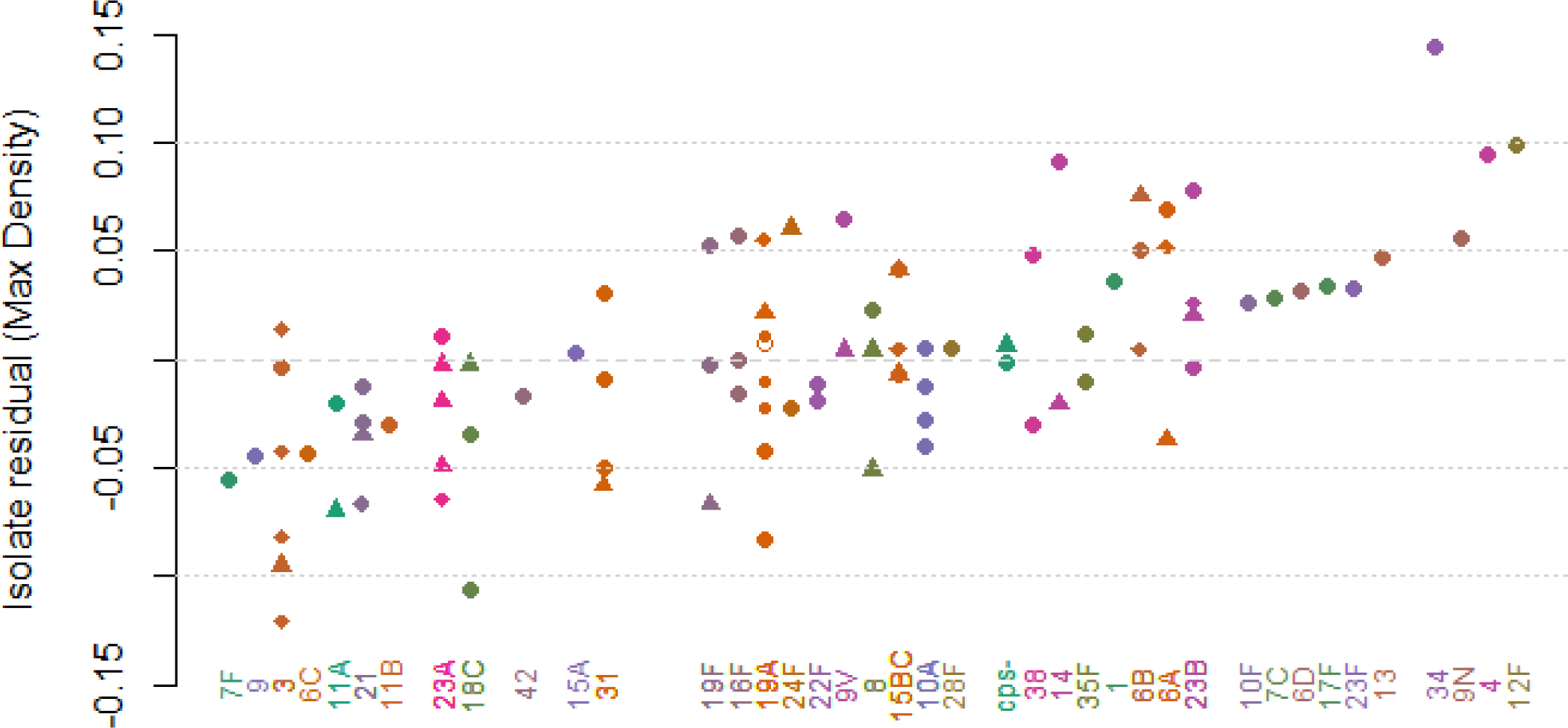
Isolate-specific random effects, organized by serotypes and genetic lineage. The random effect indicates how far above or below a strain is in maximum density achieved, on average, after adjusting for temperature, site of isolation, and presence of oxygen. Serotype is not adjusted for in this model. Each dot represents an individual isolate; within a serotype dots of the same shape share an MLST type. Only isolates for which we have sequence data are included in the plot.

**Supplementary Figures 3-6.**
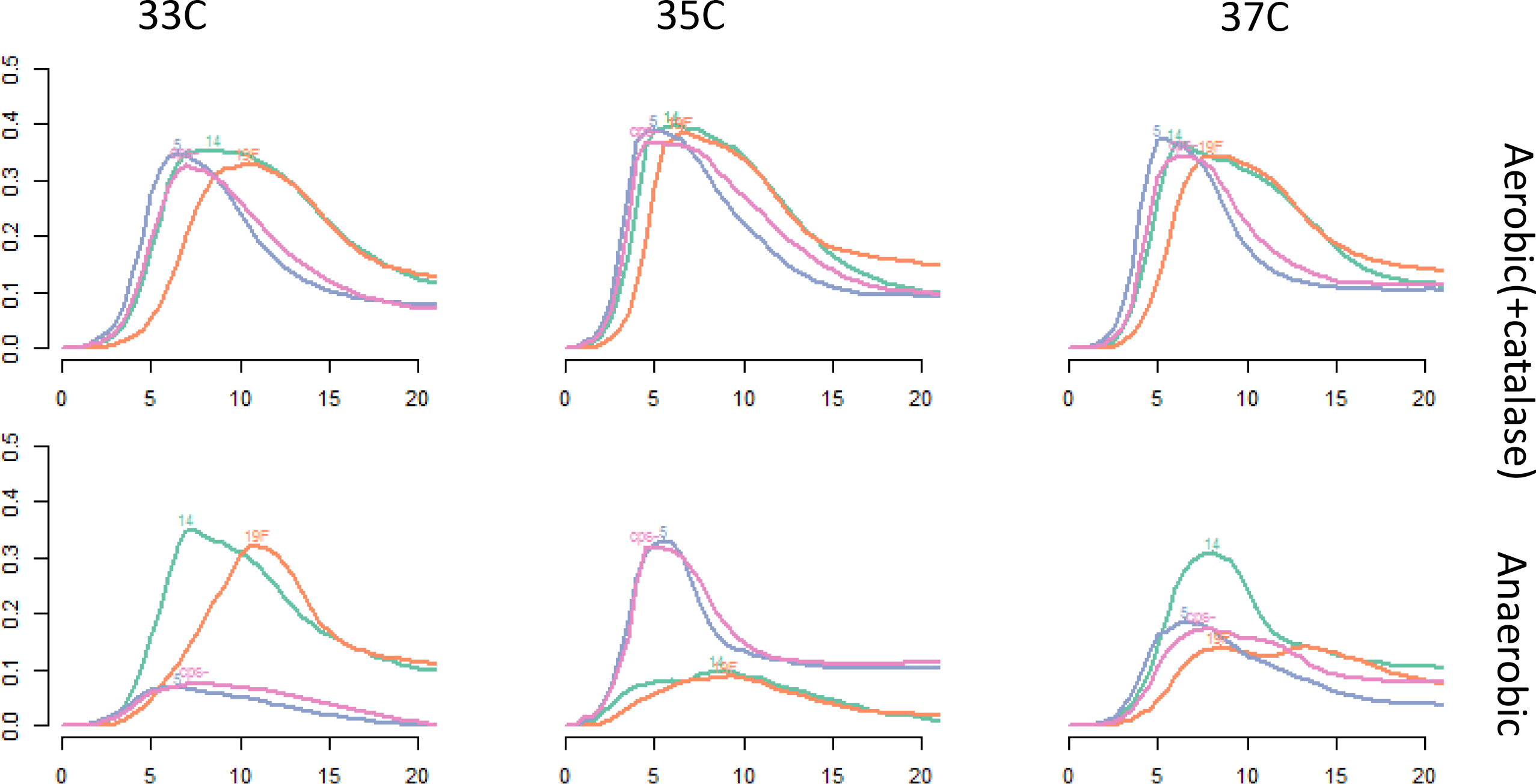
Growth curves for capsule-switch variants on the TIGR4 background and capsule knockout strains.

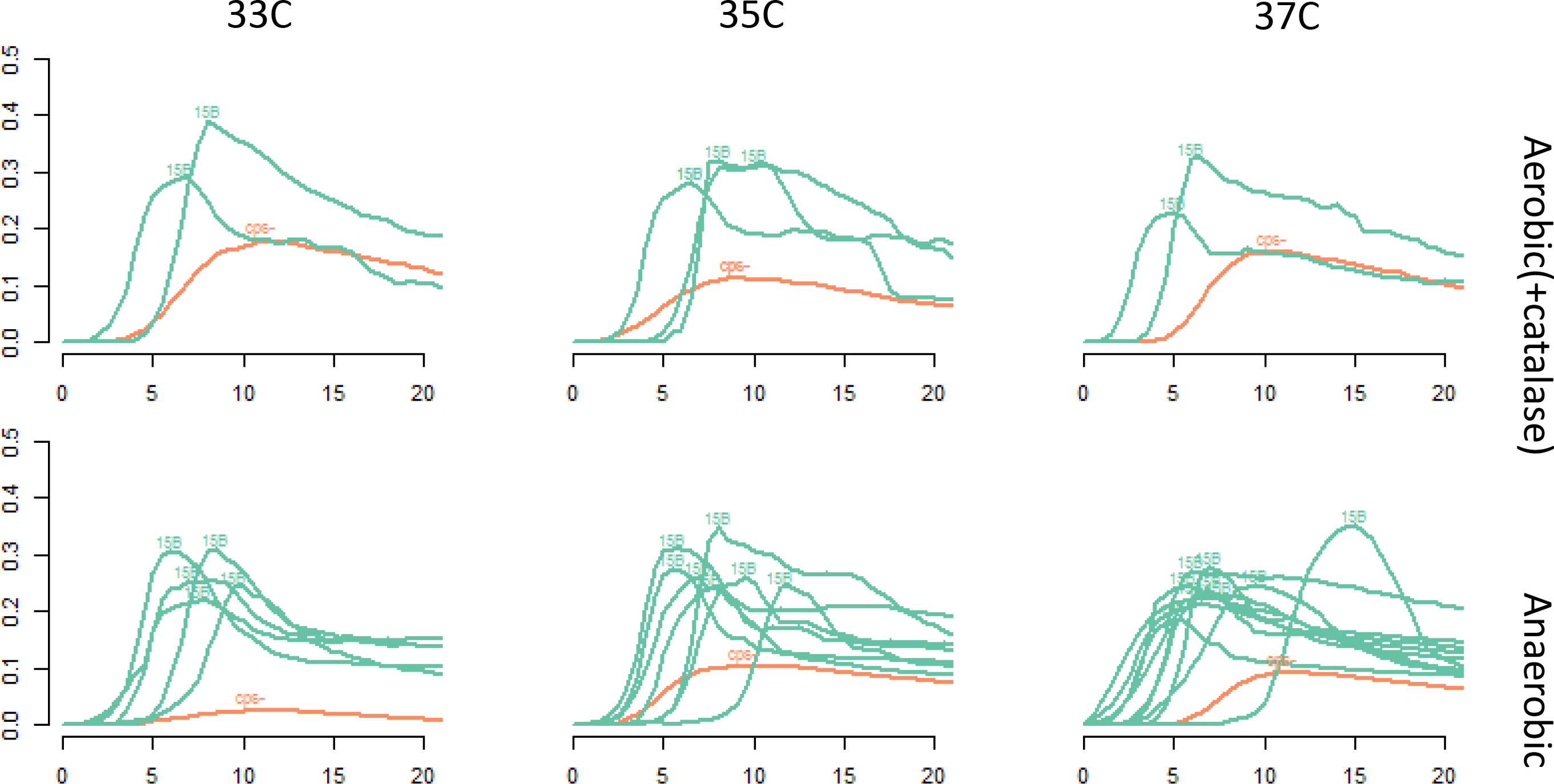

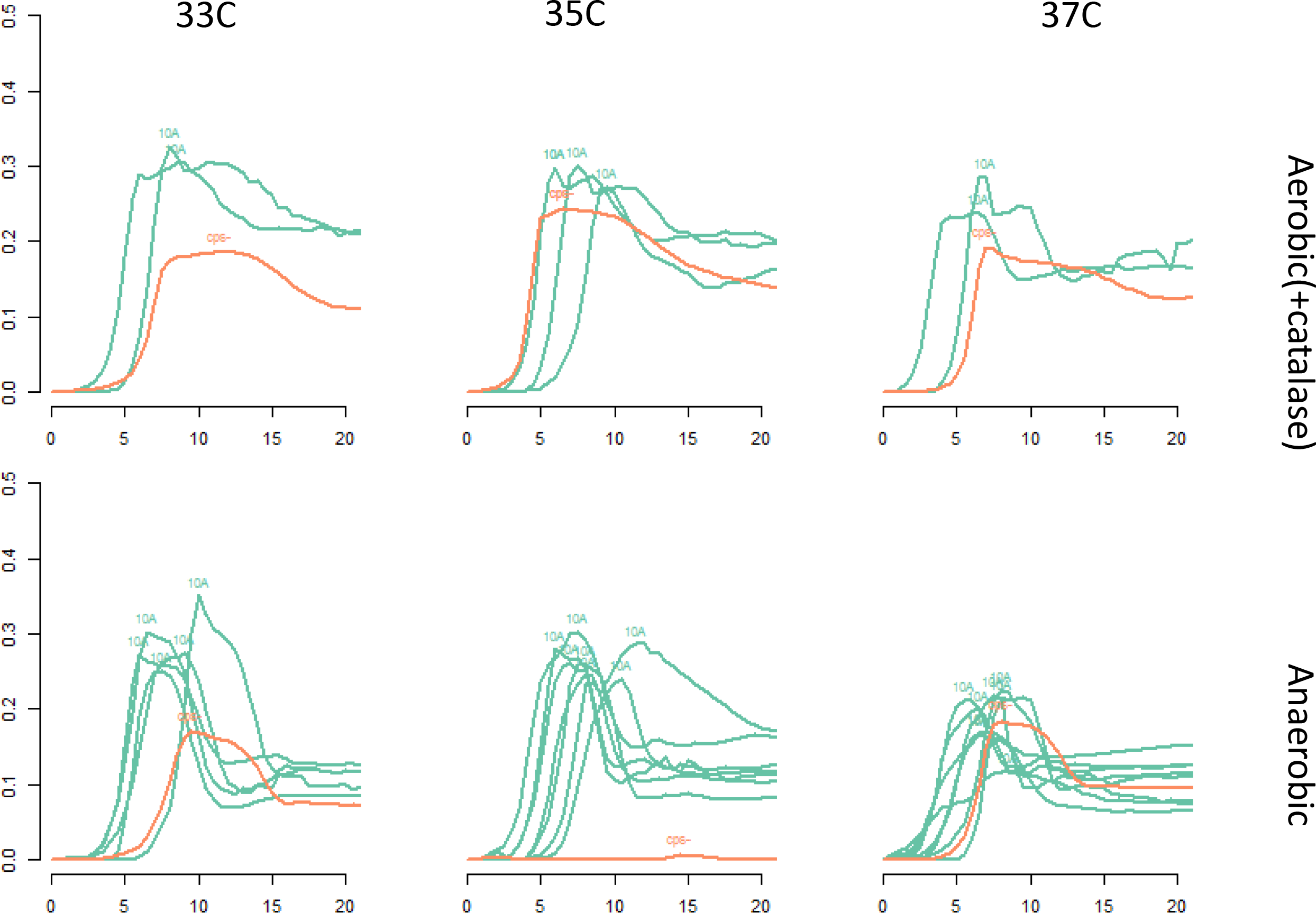

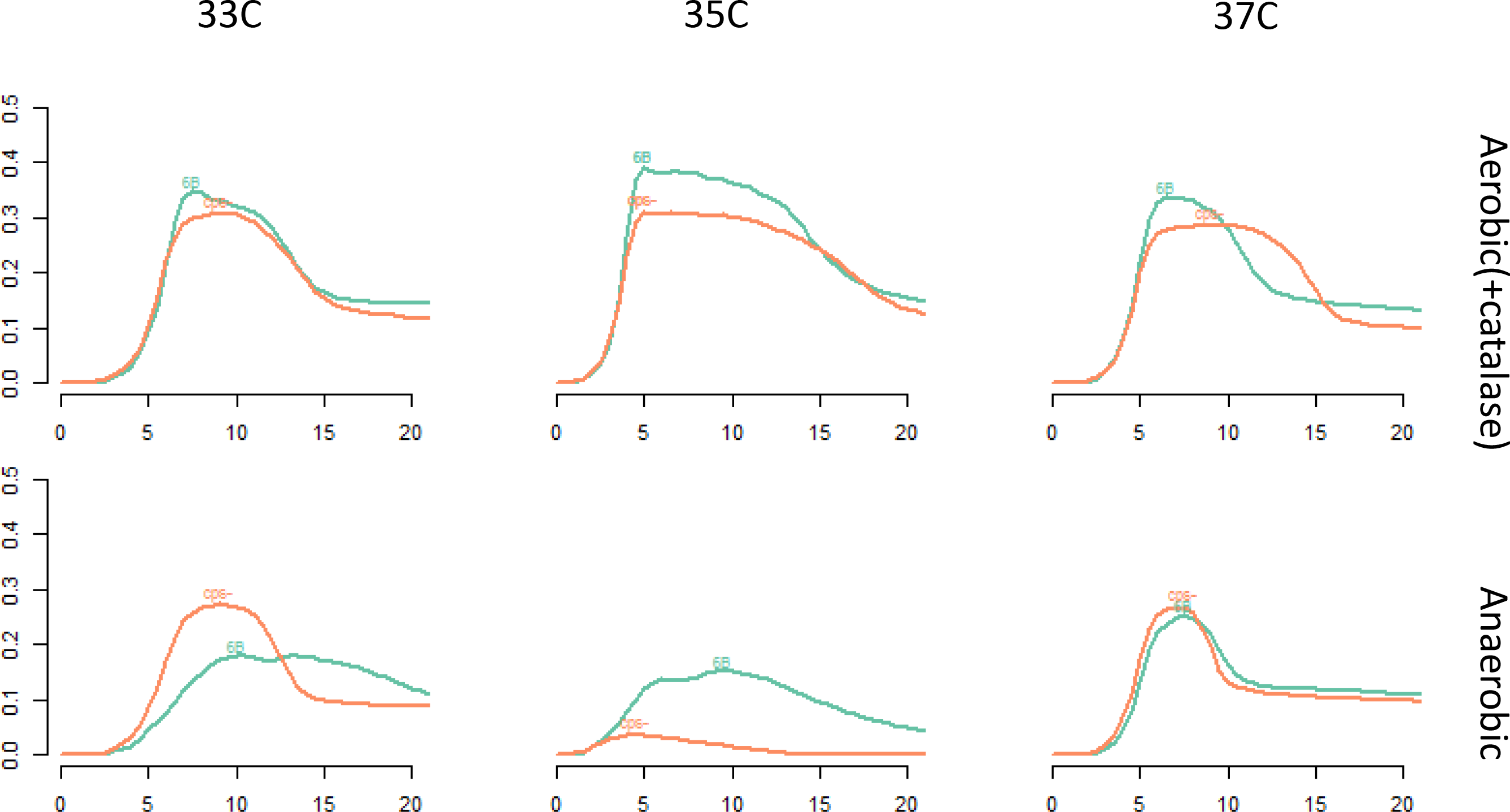

## References

1. Bogaert D, Engelen MN, Timmers-Reker AJ, Elzenaar KP, Peerbooms PG, Coutinho RA, de Groot R, Hermans PW: Pneumococcal carriage in children in The Netherlands: a molecular epidemiological study. J Clin Microbiol 2001, 39(9):3316–3320.

2. Brueggemann AB, Griffiths DT, Meats E, Peto T, Crook DW, Spratt BG: Clonal relationships between invasive and carriage Streptococcus pneumoniae and serotype- and clone-specific differences in invasive disease potential. J Infect Dis 2003, 187(9):1424–1432.

3. Geurkink N: Nasal anatomy, physiology, and function. The Journal of Allergy and Clinical Immunology 1983, 72(2):123–128.

4. Jones N: The nose and paranasal sinuses physiology and anatomy. Advanced Drug Delivery Reviews 2001, 51(1):5–19.

5. Keck T, Leiacker R, Heinrich A, Kühnemann S, Rettinger G: Humidity and temperature profile in the nasal cavity. Rhinology 2000, 38(4):167–171.

6. Van Cauwenberge P, Sys L, De Belder T, Watelet J-B: Anatomy and physiology of the nose and the paranasal sinuses. Immunology and Allergy Clinics of North America 2004, 24(1):1–17.

7. Webb P: Air Temperatures in Respiratory Tracts of Resting Subjects in Cold. Journal of Applied Physiology 1951, 4(5):378–382.

8. Ioannou S, Ebisch S, Aureli T, Bafunno D, Ioannides HA, Cardone D, Manini B, Romani GL, Gallese V, Merla A: The Autonomic Signature of Guilt in Children: A Thermal Infrared Imaging Study. PLoS ONE 2013, 8(11).

9. Keck T, Leiacker R, Riechelmann H, Rettinger G: Temperature profile in the nasal cavity. Laryngoscope 2000, 110(4):651–654.

10. Lindemann J, Keck T, Scheithauer MO, Leiacker R, Wiesmiller K: Nasal mucosal temperature in relation to nasal airflow as measured by rhinomanometry. Am J Rhinol 2007, 21(1):46–49.

11. McFadden ER, Pichurko BM, Bowman HF, Ingenito E, Burns S, Dowling N, Solway J: Thermal mapping of the airways in humans. J Appl Physiol (1985) 1985, 58(2):564–570.

12. Willatt DJ: Continuous infrared thermometry of the nasal mucosa. Rhinology 1993, 31(2):63–67.

13. Borriello G, Werner E, Roe F, Kim AM, Ehrlich GD, Stewart PS: Oxygen Limitation Contributes to Antibiotic Tolerance of Pseudomonas aeruginosa in Biofilms. Antimicrobial Agents and Chemotherapy 2004, 48(7):2659–2664.

14. Harell M, Mover-Lev H, Levy D, Sade J: Gas composition of the human nose and nasopharyngeal space. Acta Otolaryngol 1996, 116(1):82–84.

15. Hergils L, Magnuson B: Middle ear gas composition in pathologic conditions: mass spectrometry in otitis media with effusion and atelectasis. Ann Otol Rhinol Laryngol 1997, 106(9):743–745.

16. Yesilkaya H, Andisi VF, Andrew PW, Bijlsma JJ: Streptococcus pneumoniae and reactive oxygen species: an unusual approach to living with radicals. Trends Microbiol 2013, 21(4):187–195.

17. Nagaoka K, Yamashita Y, Kimura H, Suzuki M, Konno S, Fukumoto T, Akizawa K, Morinaga Y, Yanagihara K, Nishimura M: Effects of anaerobic culturing on pathogenicity and virulence-related gene-expression in pneumococcal pneumonia. The Journal of infectious diseases 2018.

18. Bättig P, Hathaway LJ, Hofer S, Mühlemann K: Serotype-specific invasiveness and colonization prevalence in Streptococcus pneumoniae correlate with the lag phase during in vitro growth. Microbes and Infection 2006, 8(11):2612–2617.

19. Hathaway LJ, Brugger SD, Morand B, Bangert M, Rotzetter JU, Hauser C, Graber WA, Gore S, Kadioglu A, Mühlemann K: Capsule type of Streptococcus pneumoniae determines growth phenotype. PLoS Pathog 2012, 8(3):e1002574.

20. Foxman EF, Storer JA, Fitzgerald ME, Wasik BR, Hou L, Zhao H, Turner PE, Pyle AM, Iwasaki A: Temperature-dependent innate defense against the common cold virus limits viral replication at warm temperature in mouse airway cells. Proceedings of the National Academy of Sciences of the United States of America 2015, 112(3):827–832.

21. Bakaletz LO: Immunopathogenesis of polymicrobial otitis media. Journal of Leukocyte Biology 2010, 87(2):213–222.

22. Giebink GS, Ripley ML, Wright PF: Eustachian Tube Histopathology during Experimental Influenza a Virus Infection in the Chinchilla. Annals of Otology, Rhinology & Laryngology 1987, 96(2):199–206.

23. Sadé J, Luntz M, Levy D: Middle ear gas composition and middle ear aeration. The Annals of Otology, Rhinology, and Laryngology 1995, 104(5):369–373.

24. Weiser JN, Bae D, Epino H, Gordon SB, Kapoor M, Zenewicz LA, Shchepetov M: Changes in availability of oxygen accentuate differences in capsular polysaccharide expression by phenotypic variants and clinical isolates of Streptococcus pneumoniae. Infect Immun 2001, 69(9):5430–5439.

25. Weiser JN, Austrian R, Sreenivasan PK, Masure HR: Phase variation in pneumococcal opacity: relationship between colonial morphology and nasopharyngeal colonization. Infect Immun 1994, 62(6):2582–2589.

26. Echlin H, Frank MW, Iverson A, Chang T-C, Johnson MDL, Rock CO, Rosch JW: Pyruvate Oxidase as a Critical Link between Metabolism and Capsule Biosynthesis in Streptococcus pneumoniae. PLoS Pathogens 2016, 12(10).

27. Aprianto R, Slager J, Holsappel S, Veening J-W: High-resolution analysis of the pneumococcal transcriptome under a wide range of infection-relevant conditions. bioRxiv 2018:283739.

28. Tóthpál A, Kardos S, Laub K, Nagy K, Tirczka T, van der Linden M, Dobay O: Radical serotype rearrangement of carried pneumococci in the first 3 years after intensive vaccination started in Hungary. European journal of pediatrics 2015, 174(3):373–381.

29. Tóthpál A, Laub K, Kardos S, Tirczka T, Kocsis A, Linden MVD, Dobay O: Epidemiological analysis of pneumococcal serotype 19A in healthy children following PCV7 vaccination. Epidemiology & Infection 2016, 144(7):1563–1573.

30. Prevaes SMPJ, de Winter-de Groot KM, Janssens HM, de Steenhuijsen Piters WAA, Tramper-Stranders GA, Wyllie AL, Hasrat R, Tiddens HA, van Westreenen M, van der Ent CK et al: Development of the Nasopharyngeal Microbiota in Infants with Cystic Fibrosis. American Journal of Respiratory and Critical Care Medicine 2015, 193(5):504–515.

31. Trzcinski K, Thompson CM, Lipsitch M: Construction of Otherwise Isogenic Serotype 6B, 7F, 14, and 19F Capsular Variants of Streptococcus pneumoniae Strain TIGR4. Applied and Environmental Microbiology 2003, 69(12):7364–7370.

32. Li Y, Thompson CM, Lipsitch M: A modified Janus cassette (Sweet Janus) to improve allelic replacement efficiency by high-stringency negative selection in Streptococcus pneumoniae. PloS One 2014, 9(6):e100510.

33. Baym M, Kryazhimskiy S, Lieberman TD, Chung H, Desai MM, Kishony R: Inexpensive multiplexed library preparation for megabase-sized genomes. PLoS One 2015, 10(5):e0128036.

34. Inouye M, Dashnow H, Raven LA, Schultz MB, Pope BJ, Tomita T, Zobel J, Holt KE: SRST2: Rapid genomic surveillance for public health and hospital microbiology labs. Genome Med 2014, 6(11):90.

35. Bates D, Mächler M, Bolker B, Walker S: Fitting Linear Mixed-Effects Models Using lme4. Journal of Statistical Software 2015, 67(1).

36. Rosario-Martinez HD: phia: Post-Hoc Interaction Analysis. R package. In., 0.2-1 edn; 2015.

37. Sleeman KL, Griffiths D, Shackley F, Diggle L, Gupta S, Maiden MC, Moxon ER, Crook DW, Peto TE: Capsular serotype-specific attack rates and duration of carriage of Streptococcus pneumoniae in a population of children. J Infect Dis 2006, 194(5):682–688.

38. Vestrheim DF, Høiby EA, Aaberge IS, Caugant DA: Impact of a pneumococcal conjugate vaccination program on carriage among children in Norway. Clin Vaccine Immunol 2010, 17(3):325–334.

39. Huang SS, Platt R, Rifas-Shiman SL, Pelton SI, Goldmann D, Finkelstein JA: Post-PCV7 changes in colonizing pneumococcal serotypes in 16 Massachusetts communities, 2001 and 2004. Pediatrics 2005, 116(3):e408–413.

40. Syrogiannopoulos GA, Grivea IN, Tait-Kamradt A, Katopodis GD, Beratis NG, Sutcliffe J, Appelbaum PC, Davies TA: Identification of an erm(A) erythromycin resistance methylase gene in Streptococcus pneumoniae isolated in Greece. Antimicrob Agents Chemother 2001, 45(1):342–344.

